# Roles of cellular senescence in driving bone marrow adiposity in radiation- and aging-associated bone loss

**DOI:** 10.1101/2021.09.07.459232

**Authors:** Abhishek Chandra, Anthony B. Lagnado, Joshua N. Farr, Megan Schleusner, David G. Monroe, Christine Hachfeld, João F. Passos, Sundeep Khosla, Robert J. Pignolo

**Affiliations:** Department of Physiology and Biomedical Engineering, Mayo Clinic College of Medicine, Rochester, Minnesota, USA; Department of Medicine, Mayo Clinic College of Medicine, Rochester, Minnesota, USA; Robert and Arlene Kogod Center on Aging, Mayo Clinic College of Medicine, Rochester, Minnesota, USA; Division of Endocrinology, Mayo Clinic College of Medicine, Rochester, Minnesota, USA

**Keywords:** aging, radiation, senescence, bone, bone marrow adiposity

## Abstract

Osteoporosis is associated with an increase in marrow adipocytes, collectively termed bone marrow adipose tissue (BMAT). An increase in BMAT is linked with decline of mesenchymal progenitors that give rise to osteoblasts which are responsible for bone accrual. Oxidative stress-induced reactive oxygen species, DNA damage, apoptosis and cellular senescence have been associated with reduced osteoprogenitors in a reciprocal fashion to BMAT; however, a direct (causal) link between cellular senescence and BMAT is still elusive. Accumulation of senescent cells occur in naturally aged and in focally radiated bone tissue, but despite amelioration of age- and radiation-associated bone loss after senescent cell clearance, molecular events that precede BMAT accrual are largely unknown. Using a mouse model here we show by RNA-Sequencing data that BMAT-related genes were the most upregulated gene subset in radiated bones. Using focal radiation as a model to understand age-associated changes in bone, we performed a longitudinal assessment of cellular senescence and BMAT. Using qRT-PCR, RNA *in situ* hybridization and histological assessment of telomere dysfunction as a marker of senescence, we observed increased *p21* transcripts in bone lining cells, osteocytes and bone marrow cells, and elevated dysfunctional telomeres in osteocytes starting from day 1 post-radiation, without the presence of BMAT. BMAT was significantly elevated in radiated bones at day 7, confirming the qRT-PCR data in which most BMAT-related genes were elevated by day 7, and the trend continued until day 42 post-radiation. Similarly, elevation in BMAT-related genes was observed in aged bones. The senolytic cocktail of Dasatinib (D) plus Quercetin (Q) – D+Q, which clears senescent cells, reduced BMAT in aged and radiated bones. MicroRNAs (miRs) linked with senescence marker *p21* were downregulated in radiated- and aged-bones, while miR-27a, a miR that is associated with increased BMAT, was elevated both in radiated- and aged-bones. D+Q downregulated miR-27a in radiated bones at 42 days post-radiation. Overall, our study provides evidence that BMAT occurrence in oxidatively stressed bone environments, such as radiation and aging, is induced following a common pathway and is dependent on the presence of senescent cells.

## 1. Introduction

Mechanisms underlying bone deterioration during physiological and pathological conditions have been a focus of study for many years. The balance between bone forming osteoblasts and bone resorbing osteoclasts maintains normal bone coupling and marked deviation from this well-orchestrated mechanism causes either osteoporosis (due to comparatively more osteoclast function) or osteopetrosis (due to relatively increased bone formation with no or minimal bone resorption)(Abhishek Chandra, Rosenzweig, & Pignolo, 2018). In common physiological conditions, such as aging or post-menopausal status, osteoporosis is the more prevalent condition and also associated with increased marrow fat or bone marrow adipose tissue (BMAT)(Veldhuis-Vlug & Rosen, 2018). BMAT in humans has been shown throughout the lifespan with no potential side effects on bone architecture up to a certain age; but with aging, disease and post-menopausal status, the presence of marrow fat appears to be inversely proportional to bone mass (Devlin & Rosen, 2015). BMAT has also been used as a predictor of bone loss showing a direct correlation to osteoporosis(Woods et al., 2020).

An increase in BMAT has been associated with depleted resident mesenchymal stem cells (MSCs) during aging and disease, as MSCs are a common precursor to both adipocytes and osteoblasts. In fact, with aging MSCs undergo linage switching toward an adipogenic fate (A. Chandra et al., 2017; Singh et al., 2016; Zhong et al., 2020). Only recently is it better understood that marrow fat cells have an autocrine, paracrine and endocrine function, with local and systemic effects(Suchacki & Cawthorn, 2018). In observations made by multiple studies, bone resorption and BMAT are directly related, but adipocyte-secreted factors that regulate bone resorption or those that potentially affect bone formation are still unknown.

Cellular senescence is a non-proliferative, biologically active state (Hayflick, 1965) that produces an active secretome known as the senescence-associated secretory phenotype (SASP)(Coppe, Desprez, Krtolica, & Campisi, 2010). Cellular senescence is one of the key underlying mechanisms accounting for age- and radiation-related bone deterioration (A. Chandra et al., 2020b; Farr et al., 2016; Farr et al., 2017). In bone tissue, senescence and SASP have been reported in osteoblasts, osteocytes, osteoprogenitors and myeloid cells, with senescent myeloid cells and osteocytes being the major contributors to the SASP(Farr et al., 2016). We have shown that BMAT is correlated with the presence of DNA damage, and mitigation of DNA damage can reduce BMAT and improve bone architecture post-radiation (A. Chandra et al., 2017; A. Chandra et al., 2018). We have also shown that focal radiation emulates several age-associated phenotypes (A. Chandra, Park, & Pignolo, 2019) with an increase in markers of senescence and SASP in bone marrow cells, osteoblasts and osteocytes (A. Chandra et al., 2020b). Interestingly, we have shown that *p21* expression was more robust at earlier time-points post-radiation than *p16*^*Ink4a*^, while *p16*^*Ink4a*^ peaked much later (A. Chandra et al., 2020b). We have previously observed a reduction in BMAT following clearance of senescent cells in aged- and radiated-bones (A. Chandra et al., 2020a; Farr et al., 2017). However, a comprehensive molecular comparison between radiation-induced BMAT and aging-induced BMAT has not yet been described.

The spectrum of adipokines that increase with cellular senescence and have the potential to cause detrimental systemic effects, are still unknown. To understand the triggers for accumulation of BMAT during aging and potential adipokine-associated genes, we used a radiation model to perform a longitudinal gene expression analysis of BMAT-related genes and compared them with those of bone tissue from old mice. We also explored the sequence of events early after radiation, to understand whether senescence precedes the accumulation of BMAT. Finally, we describe the relationships among BMAT, microRNAs (miRs) that regulate adiposity, and cellular senescence in *in vivo* models of focal radiation to bone, age-related bone loss, and pharmacologic clearance of senescent cells.

## 2. Results

### 2.1 Gene expression of adipocyte related genes in radiated bones

RNA isolated from a focally radiated 5mm area of the femoral metaphysis 3 weeks post-radiation was processed for RNA sequencing. Identified genes were then arranged based on their –log10 (False Discovery Rate). The genes with the highest –log10 (False Discovery Rate) value when depicted using a volcano plot showed that the senescence marker *Cdkn1a (p21)* and several adiposity related genes (*Cidec, Plin1, Plin4, Lpl, Igf2)* were among the most upregulated genes, while genes that were associated with endothelial function (*Vwf4, Ppbp, Pf4* and *Mmrn1)* were among the most downregulated (Fig.1A). When the genes were plotted using fold-change, we observed the same trend, and several more genes related to adipocyte function were sorted into the top 50 most up-regulated genes (Fig.1B).

**Figure 1.**
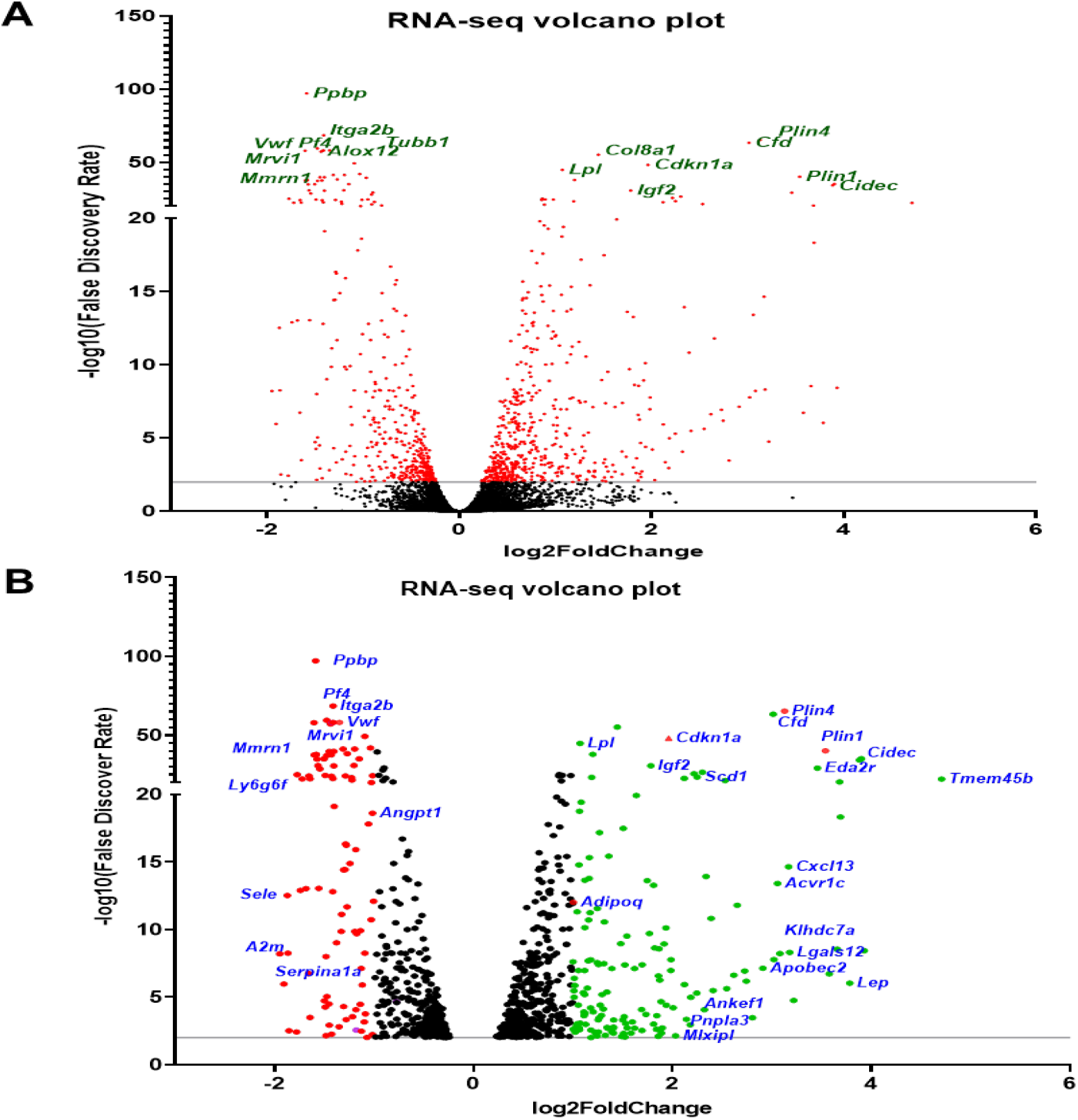
Volcano plot of the transcriptome in the radiated bones. (A) The right femoral metaphyses of C57BL/6 mice (n=3) were radiated (24Gy) in a 5 mm region, while the equivalent region of the left leg served as control. R- and NR-femurs at day 21 post-radiation were harvested for mRNA isolation. Volcano plots were generated to present the RNA-Seq data. Volcano plots of differentially expressed genes were generated from two-tailed Student’s t-test. The horizontal line and vertical lines indicate the significance threshold (FDR < 0.05) and two-fold change threshold (|log2FoldChange|>1), respectively. The differentially expressed genes (DEGs) are shown with red dots while non-DEGs are in black. (B) Upregulated DEGs with a log2Fold Change of 1 or more in the radiated bones are depicted with green-colored spots and downregulated DEGs with a log2Fold Change of 1 or more are depicted with red-colored spots. Black dots indicate all significant changes in the DEGs.

Within the gene set significantly expressed in our RNA sequencing data (1405 genes), we next curated a list of genes that play a role during adipocyte differentiation in different cell types (Table 1). Based on this curated list, we generated a heat map (Fig.2A) which showed that most of these genes were elevated in the R-bones. Based on the genes that showed close interactions using a highly stringent STRING network (Fig.2B) we used qRT-PCR to further confirm the expression of the selected genes during adipocyte differentiation of human MSCs (Supp. Fig.1).

**Figure 2.**
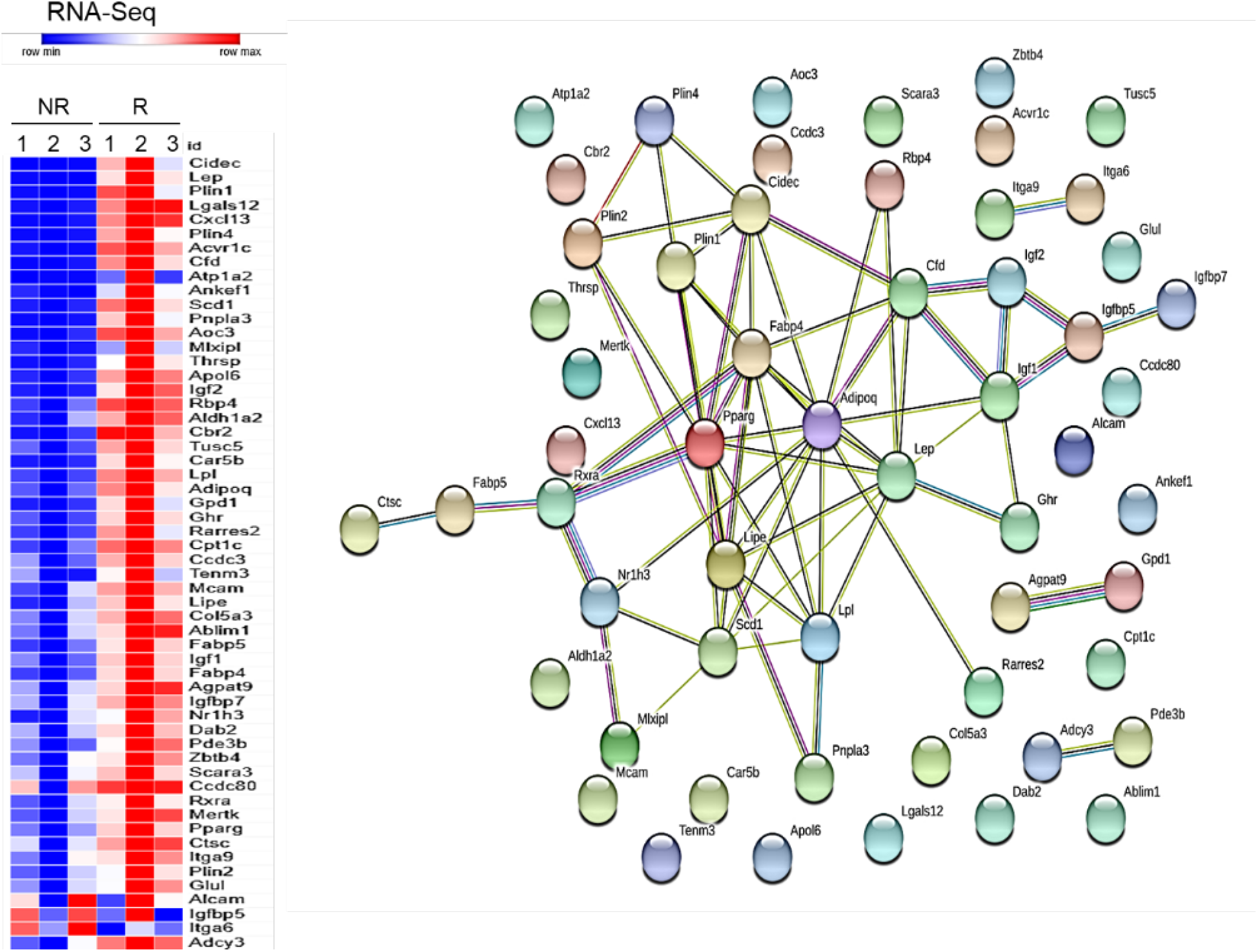
Differentially regulated adiposity related genes in radiated bones. (A) The right femoral metaphyses of C57BL/6 mice (n=3) were radiated (24Gy) in a 5 mm region, while the equivalent region of the left leg served as control. R- and NR-femurs at day 21 post-radiation were harvested for mRNA isolation. (A) A curated list of genes that are reported to be expressed during adipocyte cell differentiation was used to generate a heat map of genes expressed in R- and NR-bones detected by RNA-Seq. (B) A highly stringent STRING network representation of the BMAT-genes based on the curated heat map in (A), used for further validation by qRT-PCR.

**Table 1.**
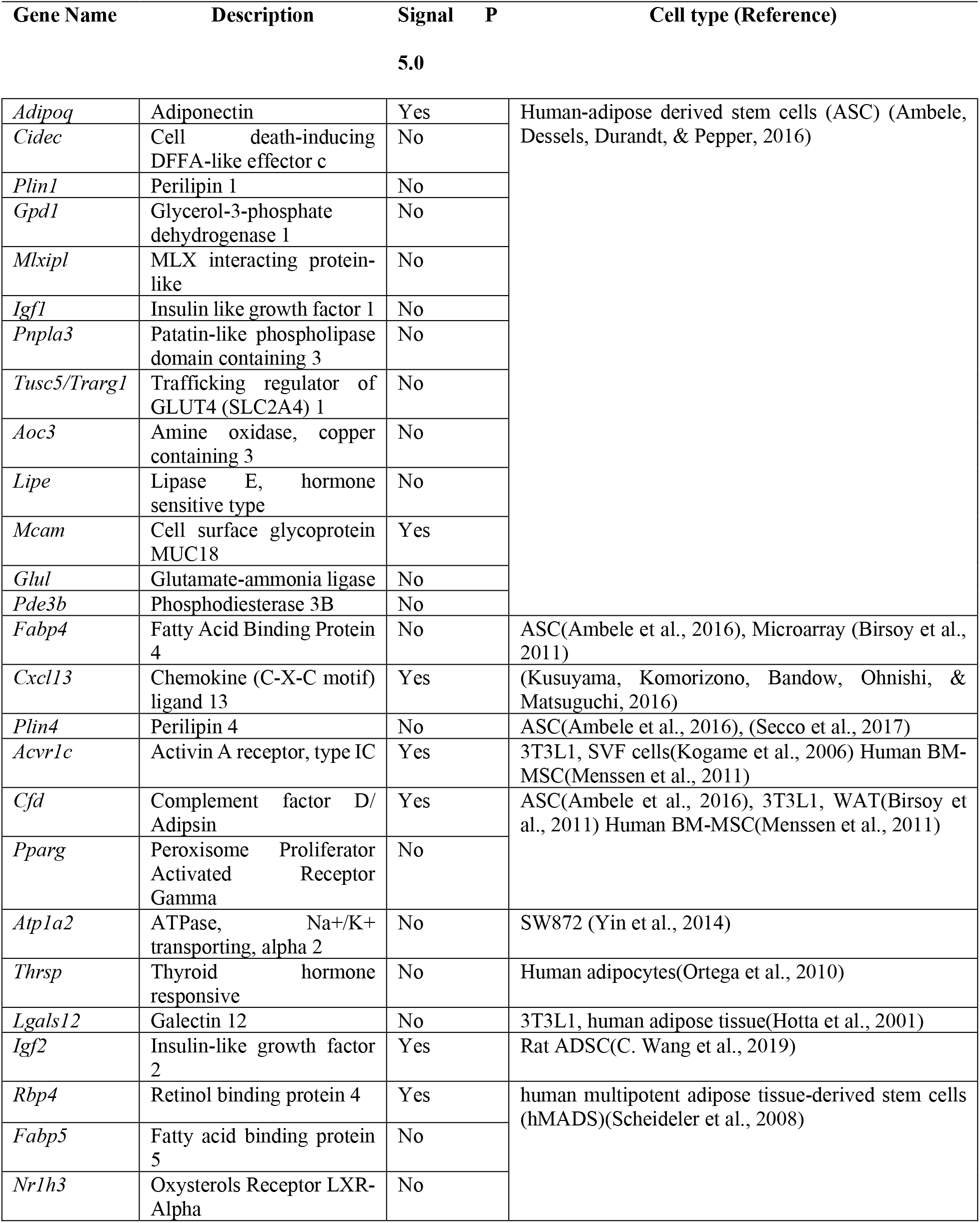
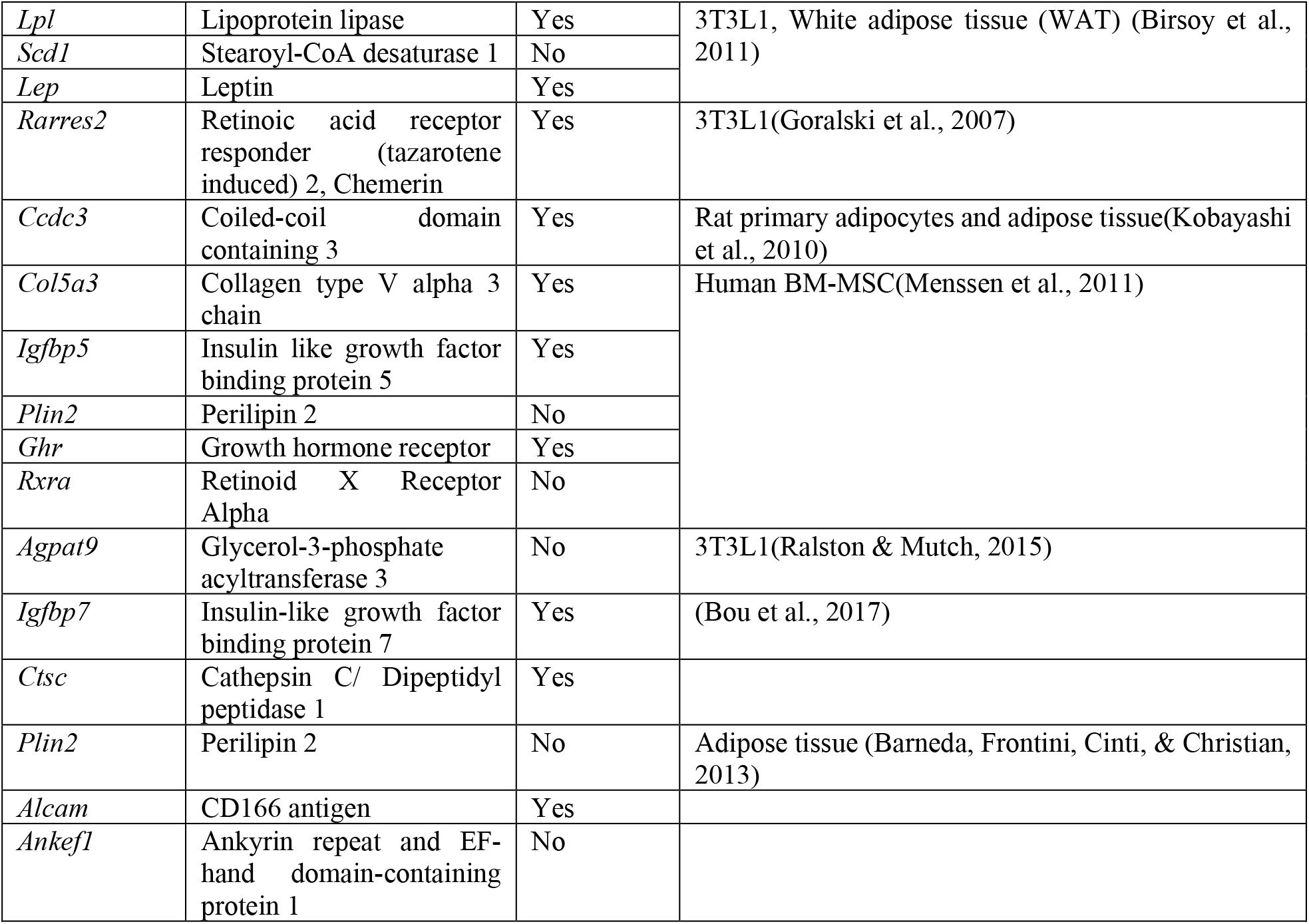
List of genes related to adipocyte function.

### 2.2 Longitudinal assessment of adiposity-related genes in radiated mouse bone tissue

To confirm the longitudinal expression of BMAT-related genes post-radiation, we analyzed 20 selected genes (as shown in Fig.2B) on days 1, 7, 21 and 42 post-radiation. We observed a non-significant elevation in the transcriptional regulators, *Pparg* and *Rxra* on day1, followed by a sustained significant increase in both these genes on days 7, 21 and 42 (Fig. 3A).

**Figure 3.**
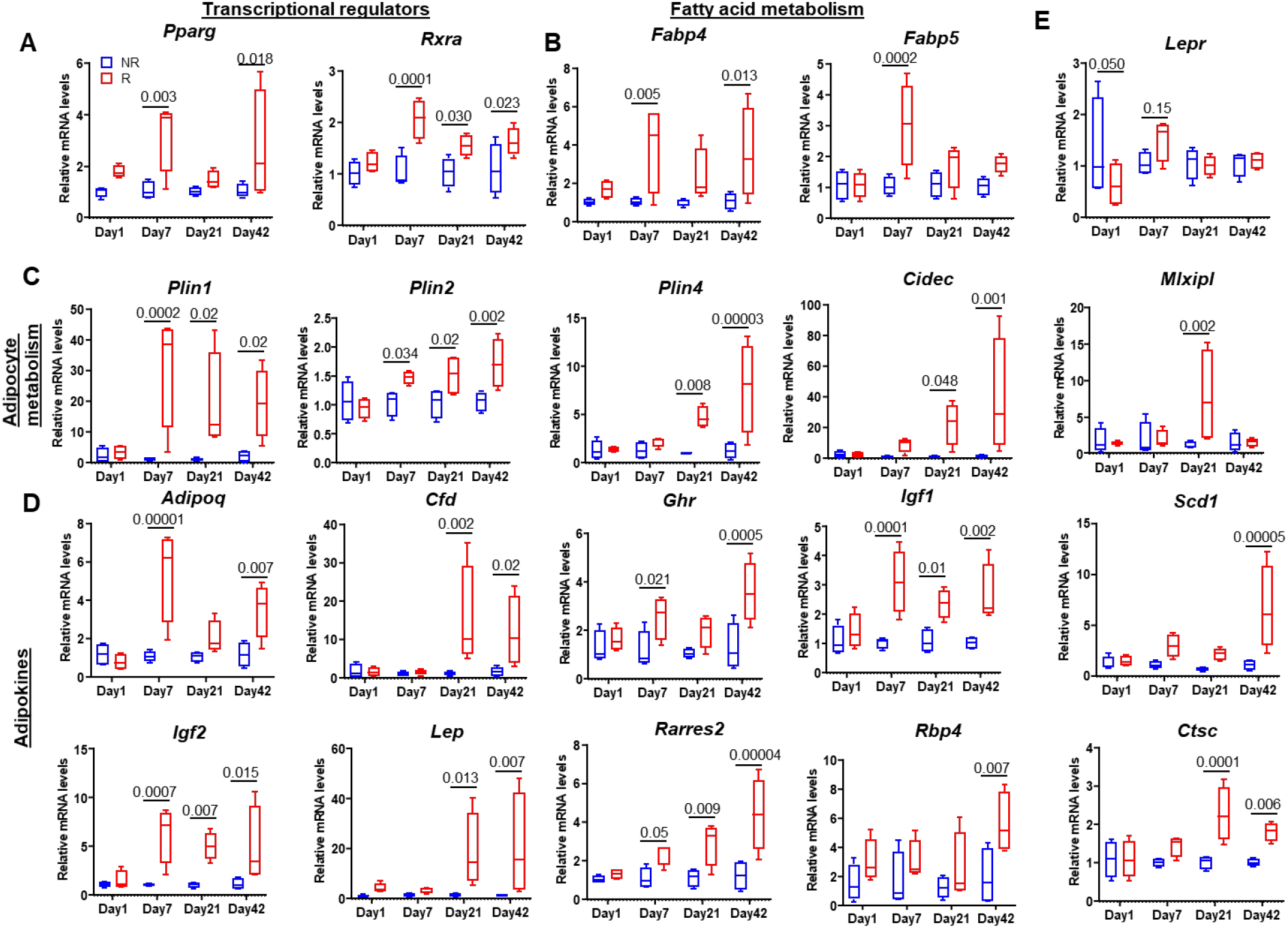
Longitudinal assessment of adiposity related genes in radiated bones. (A) The right femoral metaphyses of C57BL/6 mice were radiated (24Gy) in a 5 mm region, while the equivalent region of the left leg served as control. On days 1 (n=4),7 (n=5), 21(n=5) and 42 (n=4) R- and NR-femurs were harvested for mRNA isolation. qRT-PCR analysis is shown for genes involved with (A) transcription regulation of adipogenesis, (B) fatty acid metabolism, (C) adipocyte metabolism, (D) adipokines and (E) genes with miscellaneous functions during adipocyte differentiation. Results are expressed as medians with interquartile range. Statistical analyses were done using GraphPad Prism and p-values were calculated using a two-tailed, t-paired test.

None of the other analyzed genes were significantly expressed on day 1 post-radiation. Significant up-regulation in adipose-associated genes related to metabolism and secretory adipokines was observed in R-bones at day 7 (*Adipoq, Fabp4, Fabp5, Ghr, Igf-*1 and −2, *Plin-1* and *-2* and *Rarres2)*, day 21 (*Cfd, Ctsc, Igf-1 and −2, Lep, Mlxipl, Plin-1, −2* and *-4* and *Rarres2*) and at day 42 (*Adipoq, Cfd, Cidec, Ctsc, Fabp4, Ghr, Igf-1* and *-2, Lep, Plin-1, −2* and*-4, Rarres2, Rbp4* and *Scd1*) (Fig. 3 B-E). There were some genes which showed a non-significant elevation in R-bones at day 7 (*Cidec, Lepr* and *Scd1*) and at day 21 (*Adipoq, Cidec, Fabp4, Ghr, Rbp4* and *Scd1*).

### 2.4 Adipose-associated gene expression in old bone tissue

To understand the expression profile of adipose-related genes during aging, we next performed qRT-PCR on 24m old bones. Eighteen genes related to adipocyte transcriptional regulation, metabolism and secretory adipokines (*Adipoq, Cfd, Cidec, Ctsc, Fabp4, Ghr, Igf-1* and *-2, Lep, Lepr, Mlxipl, Plin-1, −2* and*-4, Pparg, Rarres2, Rbp4* and *Rxra)* (Fig. 4A-D) were significantly upregulated, as compared to young (5m old) mouse bone tissue. Unlike radiated bone, aged bone showed an ∼6-fold increase in the *Lepr* (leptin receptor) (Fig. 4E).

**Figure 4.**
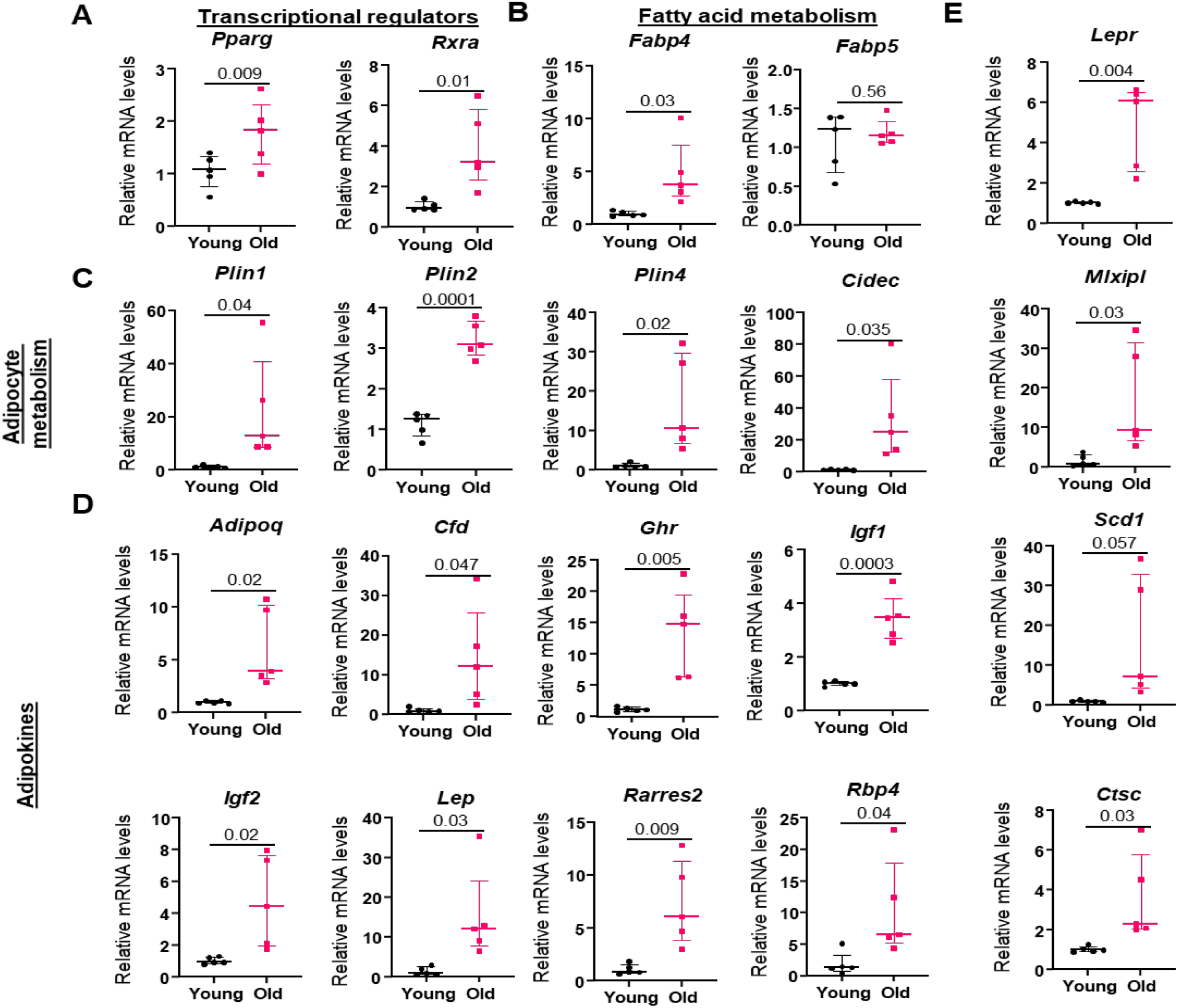
Longitudinal assessment of adiposity related genes with aging. Whole bones were collected from young (5m) (n=5) and old (24 m) (n=5) mice and mRNA were isolated. qRT-PCR analysis is shown for genes involved with (A) transcription regulation of adipogenesis, (B) fatty acid metabolism, (C) adipocyte metabolism, (D) adipokines and (E) genes with miscellaneous functions during adipocyte differentiation. Results are expressed as medians with interquartile range. Statistical analyses were done using GraphPad Prism and p-values were calculated using a two-tailed unpaired t-test.

### 2.5 Assessment of senescence markers during aging and radiation

To understand the link between senescence and adiposity seen with radiation and aging, we first confirmed our previous report that high *p21* expression was seen at early time points post-radiation, while *p16*^*Ink4a*^ expression peaked at later time points (i.e., day 42; Supp. Fig. 2A). Assessing gene expression in 24-month bone samples, we detected high *p21* (∼6-fold increase, p=0.0002) and high *p16*^*Ink4a*^ (∼3.5-fold increase, p**=**0.005) gene expression as compared to 5 month old young bone samples (Supp. Fig. 2B).

To further elucidate the dynamics of the cellular changes post-radiation and to test whether senescence precedes BMAT, we performed the TIF assay, and observed significant up-regulation in TIF^+^ osteocytes on day 1 and day 7 (Fig. 5A, B). The detection of the *p21* transcript was performed using RNA-ISH, and *p21*^*+*^-bone lining cells (BLCs), *p21*^*+*^-osteocytes (OCY) and *p21*^*+*^-bone marrow (BM) cells were detected on both days 1 and 7 post-radiation (Fig. 5C, D). Perilipin staining confirmed that adipocytes were almost absent at day 1, whereas substantial numbers of perilipin^+^ adipocytes were observed at day 7 (Fig 5E, F).

**Figure 5.**
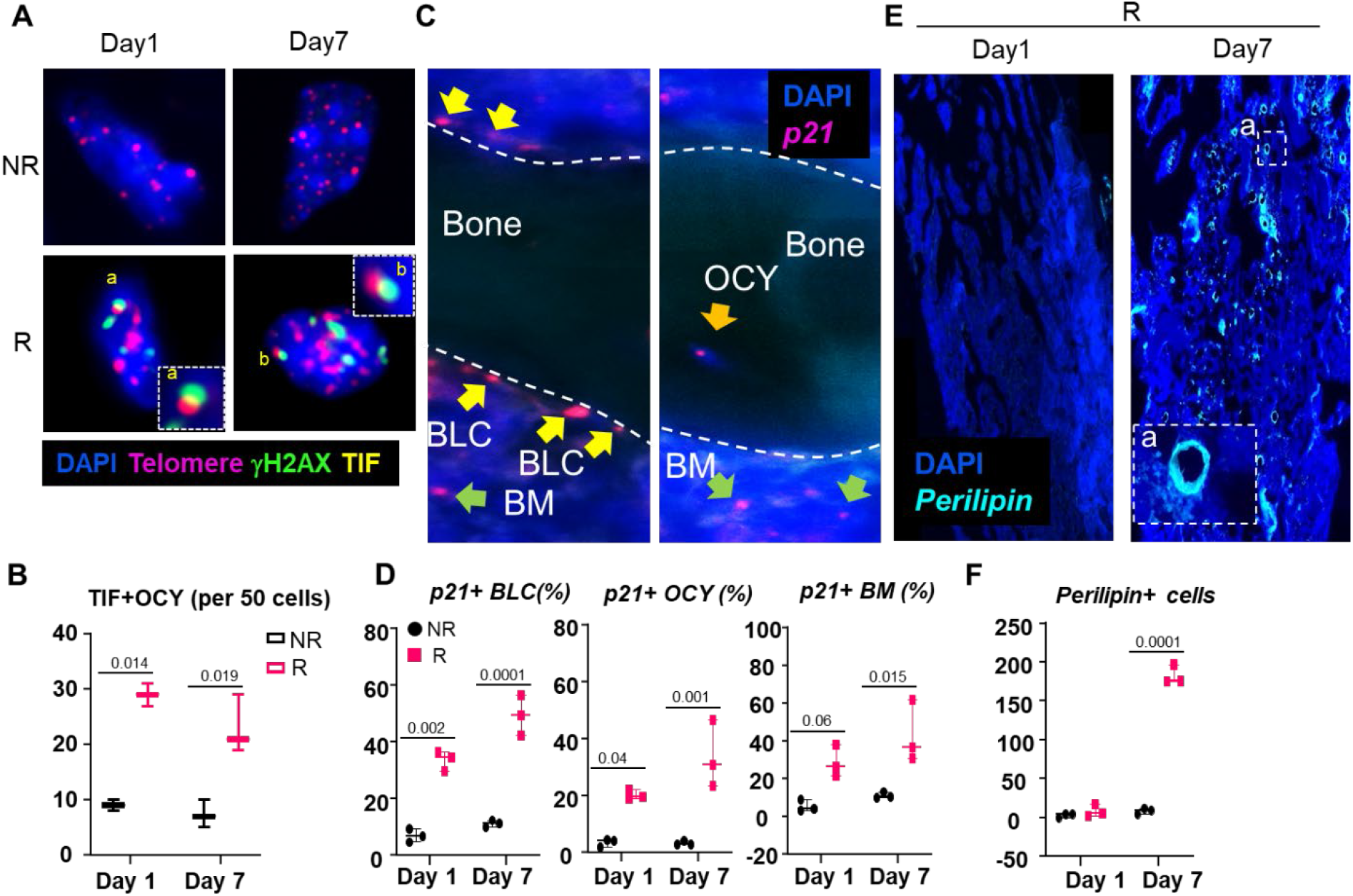
Longitudinal assessment of senescent cells and adipocytes in radiated bone tissue. To elucidate the earliest occurrence of cellular senescence post-radiation, R- and NR-bones were assessed for presence of senescent cells and adipocytes. (A, B) Colocalization of DNA damage at telomere sites, referred to as telomere dysfunction-induced foci (TIF), was visualized (A) and TIF+ osteocytes were quantified (B) at days 1 and 7 post-radiation. (C, D) RNA in-situ hybridization against Cdkn1a (p21) transcript was performed, visualized, and quantified in R- and NR-bones on days 1 and 7 and p21+ bone lining cells (BLC), osteocytes (OCY) and bone marrow (BM) cells (E, F) Perilipin stained adipocytes were assessed in NR- and R-bones. Negligible levels of adipocytes were identified in NR bones hence images are shown only for R-bones (E), while quantifications for both NR-and R-bones are presented (F). Results are expressed as medians with interquartile range. Statistical analyses were done using GraphPad Prism and p-values were calculated using a two-way ANOVA (α = 0.05) with a Tukey post hoc analysis.

### 2.6 Changes in micro-RNAs that regulate p21

Based on their binding at the 3’ UTR of *p21*, miRs were selected on their preferentially conserved targeting (PCT) as determined by the TargetScan tool which predicts the biological targets of miRNAs by searching for conserved 8mer, 7mer, and 6mer sites that match the seed region of each miRNA (Fig.6A).

**Figure 6.**
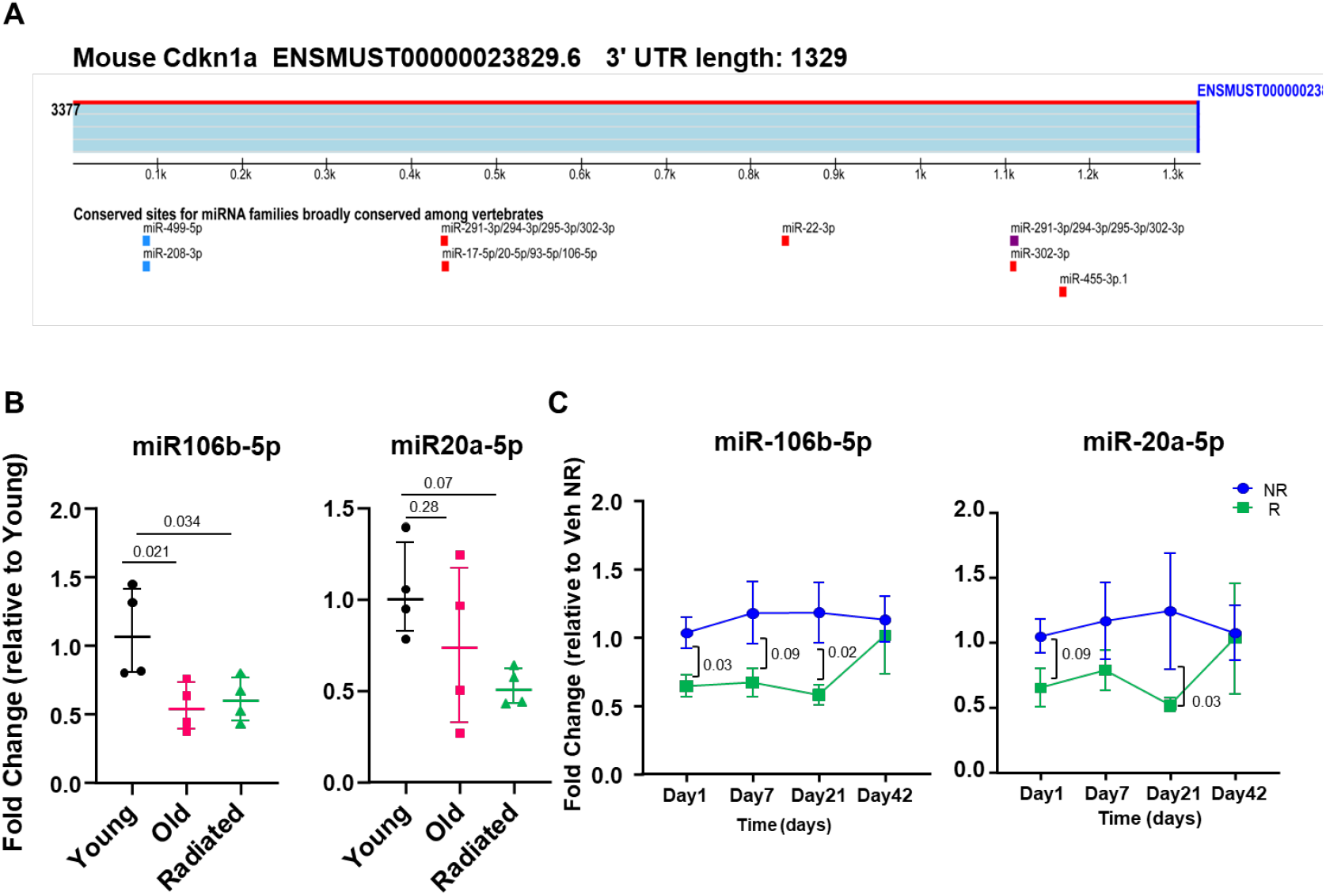
MicroRNAs targeting Cdkn1a are downregulated in radiated and aged bones. (A) A schematic showing microRNAs that bind to the 3’ UTR of the senescence marker Cdkn1a (p21). (B) Whole bones were collected from young (5m, n=4), old (24 m, n=4) and radiated (5m old, n=4) mice and mRNA were isolated. The cDNA was prepared using a miR reverse transcriptase kit and qRT PCR was performed using the primers for miR-106b-5p and miR-20a-5p. While miR-106b-5p was significantly reduced in both old- and R-bones, miR-20a-5p showed a trend to decrease in old bones but reduced significantly in radiated bones. Statistical analyses were done using GraphPad Prism and p values were calculated using a one-way ANOVA (α = 0.05) with a Bonferroni post hoc analysis. (C) To test the expression of miRs longitudinally, the right femoral metaphyses of C57BL/6 mice were radiated (24Gy) in a 5 mm region, while the equivalent region of the left leg served as control. On days 1 (n=4),7 (n=5), 21(n=5) and 42 (n=4) R- and NR-femurs were harvested for total mRNA isolation and the cDNA was prepared using a miR reverse transcriptase kit and qRT PCR was performed using the primers for miR-106b-5p and miR-20a-5p. Expression levels of both miR-106b-5p and miR-20a-5p in R-bones remained lower than NR-bones on all time points except day 42, with significant changes seen only on day 21. Results are expressed as medians with interquartile range. Statistical analyses were done using GraphPad Prism and p values were calculated using multiple t-tests.

We selected *miR-106b-5p* and *mir-20a-5p*, two miRs which have also been shown to regulate *Cdkn1a* and human aging in general (Hackl et al., 2010; Ivanovska et al., 2008). Significant down-regulation of *mir-106b-*5p (targeting *p21* mRNA) was observed both in aged-(p=0.03) and R-bones (p=0.038), while *mir-20a-5p* showed a non-significant decrease in old-bones (p=0.28), but a significant reduction in R-bones (p=0.009) when compared to young-NR bone samples (Fig. 6B). The *mir-106b-*5p was substantially reduced at different time points in the radiated bone tissue (day 1 [p=0.03], day 7 [p=0.09] and day 21 [p=0.02]), while *mir-20a-5p* was significantly down-regulated only at day 21 post-radiation (p=0.03) but had reduced expression throughout at day 1 (p=0.09) and day 7 as well (Fig. 6C).

### 2.7 Suppression of senescent cell burden reduces BMAT and BMAT-associated genes

BMAT is a common observation following radiation exposure and aging. We have showed previously that intermittent treatment with the senolytic cocktail of D+Q can suppress senescent cell burden and alleviate deleterious changes in bone architecture during aging (Farr et al., 2017) and post-focal radiation (A. Chandra et al., 2020a). Here, intermittent treatment with D+Q following 24Gy focal radiation (Fig. 7A) reduced BMAT, as measured by adipocyte number and volume, and normalized against the bone marrow area (Fig. 7B and 7C). Similarly, intermittent treatment with D+Q (Fig. 7D), suppressed bone marrow fat seen in aged bones (Fig. 7D, E), suggesting a direct role of the senescent microenvironment in promoting marrow adiposity.

**Figure 7.**
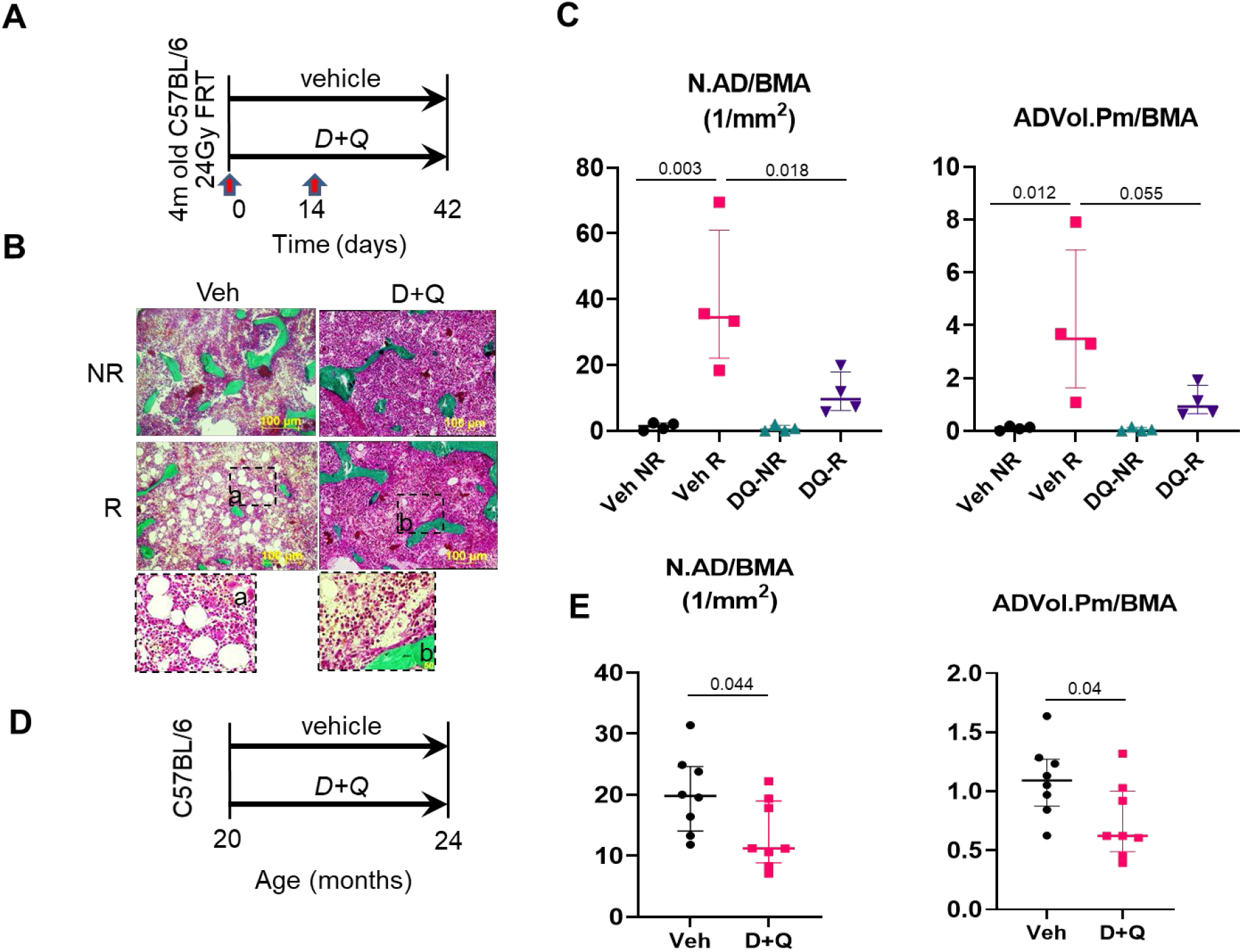
Suppression of senescent cell burden ameliorates bone marrow adiposity in radiated and aged bone tissues. (A) A schematic representing the experimental design in which right femoral metaphyses of C57BL/6 mice were radiated (24Gy) in a 5 mm region, while the equivalent region of the left leg served as control. Animals were treated with either vehicle (n=4) or D+Q (n=4) on days 0 and 14 post radiation and bones were analyzed histologically on day 42. (B) Representative images from plastic embedded 5um bone sections stained with Goldner’s trichrome stain are shown. (C) Quantification of adipocyte number and adipocyte volume normalized against the bone marrow area (BMA). Results are expressed as medians with interquartile range. Statistical analyses were done using GraphPad Prism and p-values were calculated using a two-way ANOVA (α = 0.05) with Dunnett’s post hoc analysis. (D) Schematic representation of experimental design in which 20-month old C57BL/6 received either vehicle or D+Q once a month for 4-months, and afterward the bones were processed for histomorphometry. (E) Adipocyte histomorphometric assessments are also presented for aged bones. Statistical analysis was performed using unpaired two-tailed t-test.

Although we expected that adipocyte-related genes would also be down-regulated with suppression of BMAT, we nevertheless confirmed this in our focal radiation model at day 42 post-radiation and following two doses of intermittent treatment with D+Q. We observed a robust decline in almost all adipocyte-related genes, except for *Ctsc* and the adipokines *Igf2* and *Rbp4* (Fig. 8), suggesting that these genes may be expressed by other cells as well.

**Figure 8.**
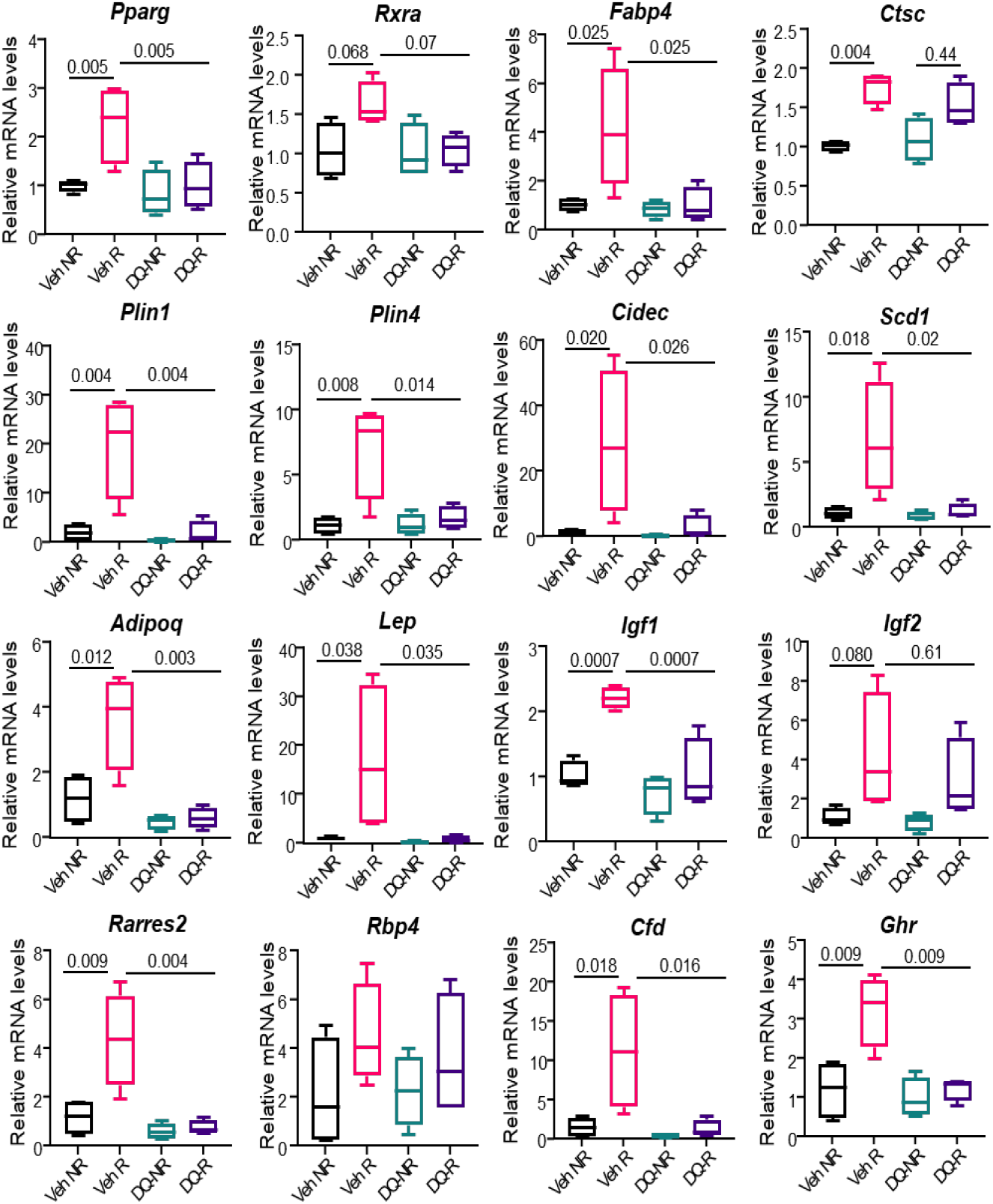
Suppression of senescent cell burden decreases BMAT-related gene expression in radiated bones. Right femoral metaphyses of C57BL/6 mice were radiated (24Gy) in a 5 mm region, while the equivalent region of the left leg served as control. Animals were treated with either vehicle (n=4) or D+Q (n=4) on days 0 and 14 post radiation and bones were harvested for mRNA. qRT-PCR was performed for 16 BMAT-related genes. Results are expressed as medians with interquartile range. Statistical analyses were done using GraphPad Prism and p values were calculated using a two-way ANOVA (α = 0.05) with a Dunnett’s post hoc analysis.

### 2.8 Adipose-related miR-27a-3p is dependent on a senescence-driven microenvironment

Intriguingly, *mir-27a-3p*, reported to be elevated during obesity and in adipose tissue as a feedback for adipose-related transcription factor expression (Lin, Gao, Alarcon, Ye, & Yun, 2009), and shown to downregulate bone marker genes, was significantly elevated in aged-(∼25-fold increase, p=0.04) and in R-bones (∼3-fold increase, p=0.017) (Fig. 9A). Up-regulation of *mir-27a-3p* was also observed in R-bones at different time points (day 1[p=0.005], day 21 [p=0.18] and day 42[p=0.04] (Fig. 9B). Treatment with the senolytic cocktail of D+Q suppressed the upregulation seen in *mir-27a-3p* levels in bones at 4 weeks post-radiation (Fig. 9C).

**Figure 9.**
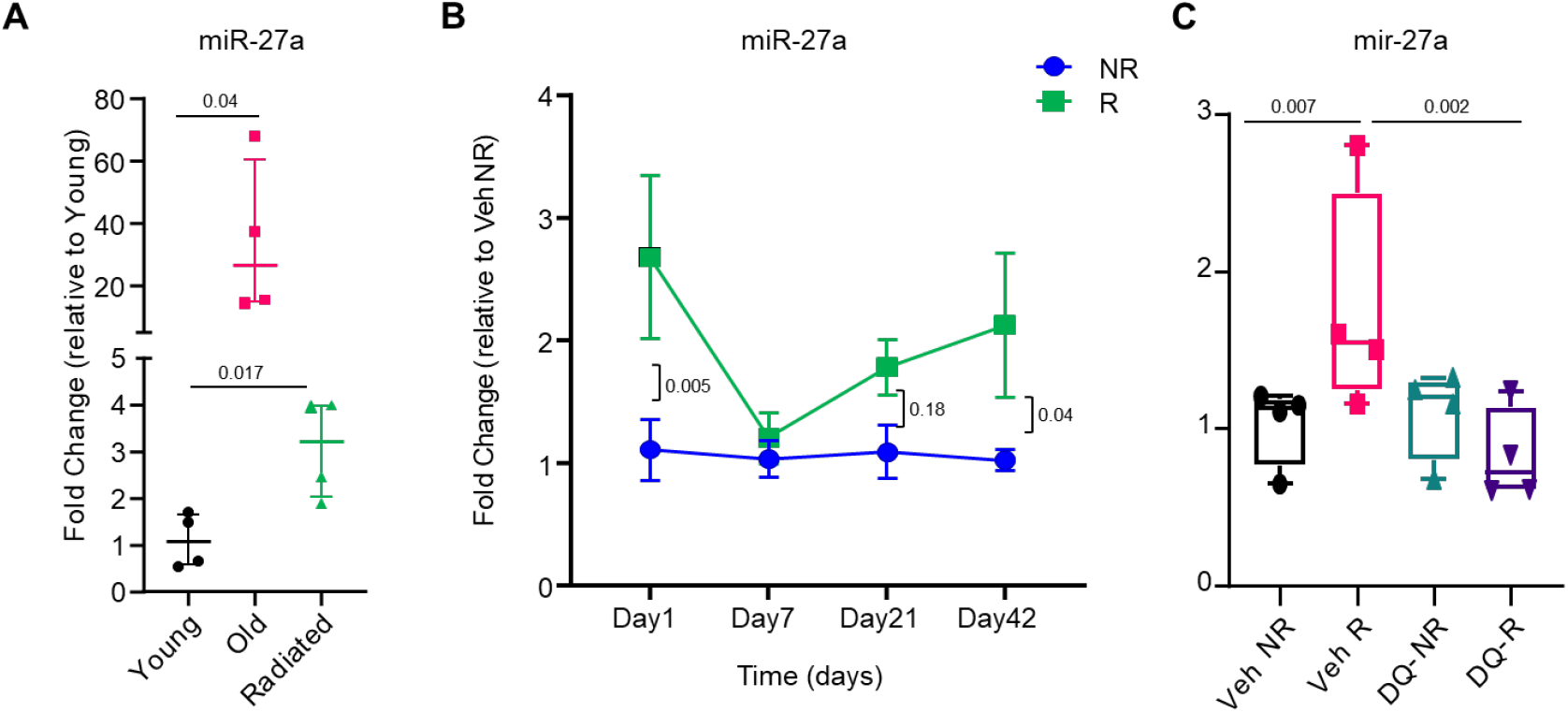
Adiposity related MicroRNA 27a is positively correlated with senescence. Whole bones were collected from young (5m), old (24 m) and radiated (5m old) mice and mRNA was isolated. The cDNA was prepared using a miR reverse transcriptase kit and qRT PCR was performed using primers for miR-27a (A). Statistical analyses were done using GraphPad Prism and p values were calculated using unpaired t-test comparing young vs old and young vs radiated. (B) The right femoral metaphyses of C57BL/6 mice were radiated (24Gy) in a 5 mm region, while the equivalent region of the left leg served as control. On days 1 (n=4),7 (n=5), 21(n=5) and 42 (n=4) R- and NR-femurs were harvested for total mRNA isolation. The cDNA was prepared using a miR reverse transcriptase kit and qRT PCR was performed using primers for miR-27a. Results are expressed as medians with interquartile range. Statistical analyses were done using GraphPad Prism and p values were calculated using a multiple-t-test. (C) Right femoral metaphyses of C57BL/6 mice were radiated (24Gy) in a 5 mm region, while the equivalent region from the left leg served as control. Animals were treated with either vehicle (n=4) or D+Q (n=4) on days 0 and 14 post radiation and bones were harvested for mRNA. The cDNA was prepared using a miR reverse transcriptase kit and qRT PCR was performed using primers for miR-27a. Results are expressed as medians with interquartile range. Statistical analyses were done using GraphPad Prism and p values were calculated using a two-way ANOVA (α = 0.05) with a Dunnett’s post hoc analysis.

## 3. Discussion

Bone is a metabolically active tissue with numerous cell types and a complex interplay of pathways that maintain an intricate balance of bone formation and resorption. Mesenchymal progenitors which generally contribute to bone formation also differentiate into marrow adipocytes, thereby causing a relative reduction in the osteoblast progenitor pool and disrupting bone homeostasis under conditions where adipogenesis is favored over osteogenesis. Increase in BMAT is a change observed across all instances of osteoporosis, is associated with reduced bone quality and fractures, and has been used as a predictor of osteoporosis (Milisic, Vegar-Zubovic, & Valjevac, 2020; Woods et al., 2020). Our previous studies support the idea that increased BMAT depletes the limited MSC pool and that MSCs undergo lineage switching toward an adipogenic cell fate in an aging microenvironment (A. Chandra et al., 2017; Singh et al., 2016). Interestingly, it was recently shown that conditional deletion of Ppar*γ* from early mesenchymal progenitors reduced BMAT but did not ameliorate thiazolidinedione- or age-associated bone loss leading the authors to a conclusion that increasing BMAT does not contribute in age-associated bone loss (Almeida et al., 2020). In contrast, deletion of BMAT by targeting adiponectin (Adipoq) expressing marrow cells was shown to induce bone mass resulting in an osteosclerotic condition (Zou et al., 2020). Furthermore, targeted deletion of Ppar*γ* in Dermo1 expressing mesenchymal lineage cells was shown to alleviate age-induced cortical bone loss in mice(Cao et al., 2020). Overall, these studies suggest that BMAT seem to have a correlation with age-related bone loss, however the trigger for BMAT during aging is still unknown.

In this study we identify cellular senescence as an important and early trigger of MSC cell fate switching toward adipogenesis during bone aging, using longitudinal assessments of senescence and BMAT-related genes in radiated bones (a model for clinical radiotherapy and accelerated bone aging), with parallel comparisons in aging bone. We have previously described the correlation between oxidative stress-induced DNA damage, a trigger for cellular senescence, and elevated levels of BMAT, as well as a reduction in BMAT with improved bone architecture (A. Chandra et al., 2017; A. Chandra et al., 2018). Down-regulation of WRN, a RecQ-type DNA helicase and DNA-repair protein, activates adipocyte-related genes in the pre-adipocyte cell line 3T3-L1 (Turaga et al., 2009). Moreover, we have shown that an aged microenvironment influences the fate of young transplanted MSCs towards adiposity (Singh et al., 2016). In this study our RNA Seq data indicated that several BMAT-related genes were upregulated 3-weeks post-radiation. As shown in Table 1 several genes are associated with *in vitro* adipocyte differentiation, but not all are studied in the context of BMAT function *in vivo*. We showed a significant elevation of expression among these genes in R-bones as shown by RNA-Seq data (Fig.2 A) and confirmed the relevance of select few in an adipocyte differentiation assay in human MSCs (Supplementary Fig. 1). A time course in radiated bones revealed that expression levels of these BMAT-related genes varied, but became more significant after 42 days, with 17 out of the 20 tested showing significant up-regulation. Plin1 staining in R-bones suggested that Plin1+ adipocytes were significantly elevated at day 7 post-radiation, but non-existent at day1 (Fig.5 E, F). This supported the gene expression data showing 11 of 20 BMAT genes being significantly upregulated at day7 but not at day1. Similarly, 18 of the tested 20 BMAT-related genes were markedly up-regulated in 24m old bones. *Lepr* was one of the genes significantly expressed in the old but not R-bones, suggesting some slight underlying differences between aging and radiation effects. Histologically, R-bone and aged bone look identical (A. Chandra et al., 2019), but this is the first evidence to our knowledge that shows an almost mirrored expression pattern in the BMAT-related genes between the two models. To understand the advent of aging-related changes to oxidative stress related cellular senescence and BMAT, we used focal radiation as a surrogate, which allowed us to assess these changes longitudinally. Our data clearly show that senescence is established on day 1 in R-bones, shown by expression of *p21* mRNA using qRT-PCR (Supp. Fig.2 A), *p21* RNA-ISH in different cells of the bone marrow compartment (Fig.5C,D), and dysfunctional telomeres in osteocytes (Fig.5A,B). It has been shown through *in vitro* experimentation that p21 is essential during adipocyte differentiation where it not only induces cell cycle arrest but also maintains adipocyte hypertrophy (Inoue et al., 2008). Here we show that *p21* expression on day 1 precedes appearance of adipocytes on day 7 post-radiation. (A. Chandra et al., 2020b; Farr et al., 2017)We also report here that the miRs that regulate *p21, miR-106b-5p (Ivanovska et al., 2008)* and *miR-20a-5p (Hackl et al., 2010)* were both downregulated at different time points post-radiation and with aging.

Genetic clearance of *p16*^*Ink4a*^ senescent cells has been shown to reduce age-related BMAT (Farr et al., 2017).Reduction in the senescent cell burden by a senolytic cocktail of D+Q has been shown to reduce both *p21* and *p16*^*Ink4a*^ transcripts as well as the SASP (A. Chandra et al., 2020b). Pharmacological treatment with the senolytic cocktail D+Q effectively clears senescent cells and suppresses the SASP in radiated (A. Chandra et al., 2020b) and aged (Farr et al., 2017) bones. Here we report that pharmacological treatment with D+Q significantly reduced radiation- and age-related adipocyte number and volume when normalized against the bone marrow area. Even though it is assumed that BMAT related genes correlate well with adipocyte numbers, we used our radiation model and tested the BMAT related genes following senescent cell clearance by D+Q. As conceptualized in a schematic (Fig.10), p21/miR-106b/miR-20a regulation drives cellular senescence in several bone marrow cells. As shown by Inoue and colleagues (Inoue et al., 2008), regulation of p21 may regulate adipocyte/BMAT related genes and thereby regulate MSC cell fate, which is also seen in R-bones. Since, by day 42, expression levels of miR-106b or miR-20a in the R-bones returned to baseline levels seen in the controls, we could not assess the effects of D+Q on either of these miRNAs. Furthermore, suppression of the SASP by D+Q (A. Chandra et al., 2020b) reduces adipokine expression and potential secretion further regulating MSC cell fate (Fig. 10). Since BMAT accumulation is inversely proportional to the MSC population and bone mineral density (A. Chandra et al., 2017), molecular regulation of BMAT could become a potential therapeutic option to treat age- and disease-related osteoporosis.

**Figure 10.**
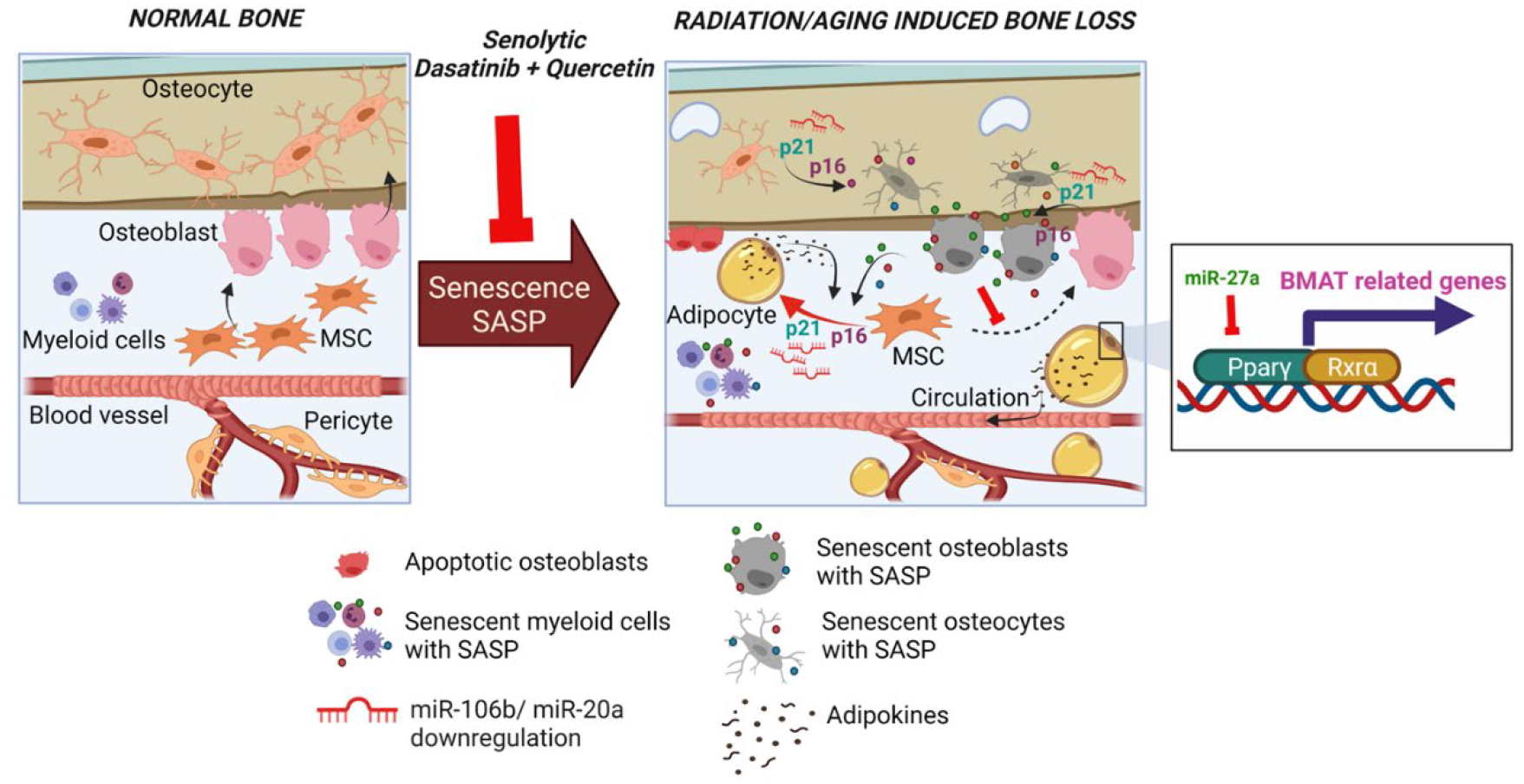
Schematic. The schematic represents changes in an oxidatively stressed, senescent microenvironment in which mesenchymal progenitors are preferentially forming adipocytes, regulated by a common process shared between aging bone and radiation-induced bone damage, and with identical expression patterns of BMAT-related genes and miRs. Increase in senescent markers p21 and p16Ink4a induce a cellular state in which production of the SASP increases and influences the bone marrow environment. Secreted adipokines can promote MSC fate switching to an adipocyte lineage. Several adipokines are also released in circulation, causing systemic effects. These changes in senescence markers and genes that regulate BMAT are in turn regulated by their corresponding miRNAs. These changes can be blocked or reversed by the clearance of senescent cells using senolytic drugs. The figure was created with BioRender.com.

To identify an upstream molecular regulator of BMAT we studied *mir-27a-3p*, a miR that regulates *Pparg* and *Rxra*, expecting for this miR to decrease. Interestingly, we observed an increase in the levels of *miR-27a-3p* both in R- and aged-bones. However, this was consistent with several reports showing elevation of *miR-27a-3p* seen during adipocyte differentiation or in adipose tissue (Yu et al., 2018). Since the primary target of *miR-27a-3p* is *Pparg* and *Rxra*, the increased levels of *miR-27a-3p* was thus an indicator of increased BMAT in which the miR-based downregulation of target genes occurs in synchrony. Hence, it is now understood that an elevated *miR-27a-3p* is a cellular negative feedback response to curb adipogenesis and counter *Pparg* or *Rxra (Deng et al., 2020)*. Interestingly, the senolytic cocktail of D+Q downregulated *miR-27a-3p*, and most of the radiation-induced BMAT genes, suggesting a direct correlation of senescent cells and elevation of BMAT. Interestingly, phosphodiesterase 3B (*Pde3B*), was the only BMAT related gene (Fig.2A, Table1) that was predicted to be regulated by both miR-106b/mir-20a and mir-27a-3p (TargetScanMouse7.1) with binding sites in the 3’UTR of the *Pde3b* gene. Pde3b has been shown to be indispensable during the maintenance of the inflammatory nature of the adipose tissue (Ahmad et al., 2016), and hence may play a role in the regulation of inflammatory cytokines including adipokines. Some of these cytokines which are part of the inflammasome may be traditional SASP factors, but some of them can be characterized as novel SASP factors.

Adipokines, the adipocyte derived hormones, also referred to as inflammatory cytokines secreted by the adipocytes, have detrimental effects on bone homeostasis. It is well understood now that adipokines and other secreted proteins from adipocytes interact with other cell types in the bone in a paracrine function, and potentially other organs of the body in a systemic manner (MacDougald & Burant, 2007). Adipokines are known to potentiate the formation of new adipocytes and are often inflammatory in nature. Amongst the BMAT-related genes curated by us and expressed significantly in R-bones, we found several known adipokines, but some BMAT proteins previously not described as adipokines had a secretory signal sequence predicted by the Signal P software, suggesting that these proteins could be potential adipokines (Table1).Chemerin is one such adipokine, which has been studied extensively. Early reports showed that the *Chemerin (Rarres2)* gene and its receptor gene *CMKLR1* (chemokine like receptor-1), were expressed in several organs (Bozaoglu et al., 2007) and cell types, including pre-adipocytes, macrophages and mature adipocytes (MacDougald & Burant, 2007). In our studies *Rarres2* expression increased in R-bones in a sequential manner, while being significantly expressed in the old-bones. Chemerin has been shown to induce osteoclastogenesis, and a neutralizing antibody against Chemerin blocks this process (Muruganandan, Dranse, Rourke, McMullen, & Sinal, 2013). Some other adipokines such as Leptin (*Lep* gene), Adipsin (*Cfd* gene), Adiponectin (*Adipoq* gene), Visfatin (*Nampt* gene) and Vaspin (*Serpina12* gene) have been used as biomarkers that are inversely associated with bone health (Cervellati et al., 2016; Terzoudis et al., 2016), however detailed mechanisms are still unknown. While the *Nampt* gene was not detected in our RNA Seq data, *Serpina12* was upregulated significantly (data not shown). In this study we show that the clearance of senescent cells could regulate adipokine expression in R-bones.

Based on the longitudinal studies in R-bones and pharmacological manipulation by D+Q, our data strongly suggests that senescence is the driving mechanism for MSC cell fate conversion to adiposity. Based on the gene expression profile of aged- and R-bones, our data also suggest that changes with radiation in an acute manner may represent an accelerated aging phenotype and may be used as a surrogate for physiological age-associated bone damage. Furthermore, while suppression of SASP factors as a group could regulate BMAT and maintain MSC cell fate, targeting individual adipokines may also regulate MSC cell fate and thus become a potential therapeutic to target age- and disease-related bone loss.

## 4. Materials and Methods

### 4.1 Animal study design

All animal studies were approved by the Institutional Animal Care and Use Committee at the Mayo Clinic. The animals were purchased from The Jackson Laboratory and housed in our facility at 23–25°C with a 12-h light/dark cycle and were fed with standard laboratory pellets with free access to water. Four-month-old C57BL/6 mice received focal radiation as a single dose of 24Gy on day 0. Using X-Rad-SmART (Precision X-Ray (PXi), North Branford, Connecticut) an image-guided focal dose of 24Gy at 6.6Gy/minute was delivered to a 5mm region of the distal metaphyseal region of the right femur, while the left femur served as the contralateral control. Bones were harvested at day 1, 7, 21 and 42 for gene expression and RNA sequencing (day 21). Additional animals were radiated, and bones were collected on days 1 and 7 for histology and RNA-*in situ* hybridization (ISH). To test the effect of senolytic cocktail Dasatinib (D, 5mg/kg) and Quercetin (Q, 50mg/kg) on radiation-induced BMAT related genes, D+Q was dosed on day 0 and day 14 post-radiation as described previously, and bones were collected on day 42 post-radiation (A. Chandra et al., 2020b). To test the effect of D+Q on age-related BMAT, 20 month-old C57BL/6 mice were injected with vehicle or D+Q once monthly for 4 months as described previously (Farr et al., 2017), with assessments of BMAT performed in the vertebra.

### 4.2 RNA preparation, sequencing, and bioinformatics

A 5 mm region below the growth plate of the distal metaphyseal femur was cut out from the radiated (R) and non-radiated (NR) legs. After removal of muscle tissue, the bone samples were homogenized and total RNA was isolated using RNeasy Mini Columns (QIAGEN, Valencia, CA) for subsequent RNA-sequencing, which was performed by the Mayo Sequencing Core. MAP-RSeq version 3.0.2 (Kalari et al., 2014), an integrated RNA-Seq bioinformatics pipeline developed at the Mayo Clinic, was used for comprehensive analysis of raw RNA sequencing paired-end reads. MAP-RSeq employed STAR (Dobin et al., 2013), the very fast, accurate and splice-aware aligner for aligning reads to the reference mouse genome, build mm10. The aligned reads were then processed through multiple modules in a parallel fashion. Gene and exon expression quantification was performed using the Subread (Liao, Smyth, & Shi, 2013) package to obtain both raw and normalized (FPKM – Fragments Per Kilobase per Million mapped) reads. STAR-Fusion, a module to detect fusions in STAR (Dobin et al., 2013), was used to identify and report any expressed gene fusions in the samples. Likewise, expressed single nucleotide variants (SNVs) and small insertions-deletions (INDELs) were detected using a combination of bioinformatics tools such as GATK (McKenna et al., 2010), HaplotypeCaller(McKenna et al., 2010) and RVBoost (C. Wang et al., 2014). Finally, comprehensive quality control modules from the RSeQC(L. Wang, Wang, & Li, 2012) package were run on the aligned reads to assess the quality of the sequenced libraries. Results from these modules described above were linked through a single html document and reported by MAP-RSeq.

R bioinformatics package DESeq2 (Love, Huber, & Anders, 2014) was used for differential gene expression analysis. The criteria for selection of significant differentially expressed genes were p-adjusted ≤ 0.05 and | log2 fold change | ≥ 1.5. The RNA-Seq data has been uploaded on the GEO database (https://www.ncbi.nlm.nih.gov/geo/query/acc.cgi?acc=GSE180076).

### 4.3 Adipocyte differentiation assay

Human MSCs were maintained in growth media (Dulbecco’s Modified Eagle Medium (DMEM, Gibco) containing D-glucose (1g/L), L-Glutamine and sodium pyruvate (110mg/L)) supplemented with 10% fetal bovine serum (Gibco). For adipocyte differentiation, cells were cultured in growth media supplemented with indomethacin (60μM), Isobutyl methylxanthine (500μM), dexamethasone (1μM) and insulin (10μg/ml). Cells were cultured in growth media or adipocyte differentiation media for 14days. Adipocyte differentiation was confirmed by visually confirming the lipid droplet accumulation by oil red staining (Supp. Fig.1A). From a parallel culture mRNA was collected from control (n=2) and adipogenic differentiated cells (n=2) for gene expression analysis.

### 4.4 Telomere fluorescence *in situ* hybridization (FISH) and DNA damage

Paraffin embedded bone sections were de-paraffinized followed by serial hydration in 100% and 70% ethanol, with final hydration in PBS. Antigen was retrieved by incubation in 0.01 M citrate buffer (pH 6.0) at 95 °C for 15 min, placed on ice for 15 min and then washed in water and PBS for 5 min each. Subsequently, blocking buffer was applied (normal goat serum 1:60 in PBS/BSA, #S-1000; Vector Laboratories) for 30min at room temperature (RT) followed by primary antibody (γ-H2A.X, Cell Signaling, mAb #9718) overnight at 4 °C. Slides were washed three times with PBS and incubated for 60 min with a species-specific secondary antibody (no. PK-6101; Vector Lab). Sections were then washed 3 times in PBS for 5 min and Cy5 Streptavidin (1:500 in PBS, No: SA-1500, Vector Lab) was applied for 25 min followed by three washes in PBS, crosslinking with 4% paraformaldehyde for 20 min and dehydration d in graded ethanol. Sections were denatured for 10 min at 80 °C in hybridization buffer [70% formamide (Sigma UK), 25 mM MgCl2, 0.1 M Tris (pH 7.2), 5% blocking reagent (Roche, Germany)] containing 2.5 μg/mlCy-3-labelled telomere-specific (CCCTAA) peptide nucleic acid probe (Panagene), followed by hybridization for 2h at RT (minimum) in a humidified chamber in the dark. Next, slides were washed once with 70% formamide in 2 × SSC for 10 min, followed by 1 wash in SSC and PBS for 10 min. Sections were mounted with Dapi (ProLong™ Diamond Antifade Mountant, Invitrogen, P36962). In-depth Z-stacking was used (a minimum of 135 optical slices using a 63X objective). Number of TIFs per cell was assessed by manual quantification of partially or fully overlapping (in the same optical slice, Z) signals from the telomere probe and γH2A.X in z-by-z analysis. Images were deconvolved with blind deconvolution in AutoQuant X3 (Media Cybernetics).

For Formalin-Fixed Paraffin-Embedded (FFPE), immunohistochemistry was carried out following an overnight incubation with rabbit monoclonal anti-perlipin1 (1:100, Cell Signaling Technology, 9349). The next day following three washes, tissues were incubated with a species-specific secondary antibody (Alexa 647) for 1h then washed 3 times with PBS and mounted using ProLong Gold Antifade Mountant with DAPI (Invitrogen).

### 4.5 RNA *in situ* hybridization (RNA-ISH) and Immunohistochemistry

RNA-ISH was performed per the RNAScope protocol from Advanced Cell Diagnostics Inc. (ACD): RNAScope Multiplex Fluorescent Assay v2. Briefly, the assay allows simultaneous visualization of up to 4 RNA targets, with each probe assigned a different channel (C1, C2, or C3 or C4). Each channel requires its own amplification steps - for example, p21 (CDKN1A) C1 probe was amplified by HRP-C1, followed by the addition of whichever fluorophore was assigned to that probe/channel using the Opal dyes (Opal 570 (Cy 3 Range), then blocked to perform the next amplification in another channel, p16/p19 (CDKN2A) C3 using Opal 650 (Cy5). Tissue sections were then mounted using ProLong Gold Antifade Mountant with DAPI (Invitrogen). Sections were imaged using in-depth Z stacking.

### 4.6 Quantitative RT-PCR

A 5 mm region below the growth plate of the distal metaphyseal femur was cut out from the radiated and the non-radiated legs. After removal of muscle tissue, the bone samples were stored in TRIzol (Invitrogen) at −80°C. Bones were homogenized and total RNA was isolated using a phase separation method, followed by a RNase-free DNase treatment to remove genomic DNA and a cleanup of samples with the RNeasy Mini Columns (QIAGEN, Valencia, CA). The quality of the RNA was determined using a Nanodrop spectrophotometer with A260/230 ≥1.6 and A260/A280 set at ≥1.8 (Thermo Scientific, Wilmington, DE). The cDNA was generated from mRNA using the High-Capacity cDNA Reverse Transcription Kit (Applied Biosystems by Life Technologies, Foster City, CA), according to the manufacturer’s instructions.

Primers which were designed for our previous studies (Farr et al., 2016) were used for RT-qPCR in this study as well. Additional primers were designed using Primer Express ® Software Version 3.0 (Applied Biosystems, Foster City, CA) (Supplementary Table S1 and S2). Using a high throughput ABI Prism 7900HT Real Time System (Applied Biosystems, Carlsbad, CA) with SYBR Green (QIAGEN, Valencia, CA) as the detection method, gene expression was detected. Gene expression was normalized against the average of 2 reference genes (Actb and Tuba1) and the fold difference between the reference gene and the target gene [reference cycle threshold (Ct)-gene of interest Ct] was calculated by comparing the median Ct with NR-femurs serving as control for R-femurs, and young bone cells serving as controls for old.

### 4.7 Statistical analysis

Data are expressed as means ± standard error (SEM) and analyzed by two-tailed paired Student’s t-test for comparison of R and NR femurs (two-tailed). Individual animals were treated as indicated in the figure legends. The R-based tool from Bioconductor, edgeR, was used to perform the differential expression analysis. Those mature mRNAs were significantly differentially expressed at a p-value < 0.05 and a log2 fold change higher than 1 or less than −1. Heat maps were created using Morpheus software(https://software.broadinstitute.org/morpheus). For D+Q studies, statistical analyses were done using GraphPad Prism and p values were calculated using a two-way ANOVA with a Dunnett’s post hoc test performed to compare vehicle-R with all the other groups. The D+Q studies done in the aging cohort were analyzed using a two-tailed unpaired t-test. Statistical comparisons of young, old, and radiated miRs were done using one-way ANOVA with a Bonferroni post-hoc test.

## Acknowledgements

This work was made possible by the Eagle’s Cancer Research Fund (to AC), Mayo Clinic Clinical and Translational Science Award (CTSA), grant number UL1TR002377, from the National Center for Advancing Translational Science (NCATS), a component of the National Institutes of Health (NIH) (to AC) and UL1TR000135, Center for Clinical and Translation Science (CCATS)(to DGM and AC), as well as the Robert and Arlene Kogod Professorship (to RJP); P01 AG062413 (SK, JNF, RJP), P01 AG004875 (SK/DGM), R01 AG048792 (SK/DGM), R01 AR068275 (DGM), and, R01 DK128552 (JNF), and K01 AR070241 (JNF).

## Author Contributions

AC and RJP designed the study, AC, AL, JNF, MS and CH performed the experiments, while AC, DM, JNF, SK, JP and RJP analyzed and interpreted the data. AC and RJP wrote the manuscript and all authors revised and approved the final version of the manuscript.

---

**Supplementary Figure 1.**
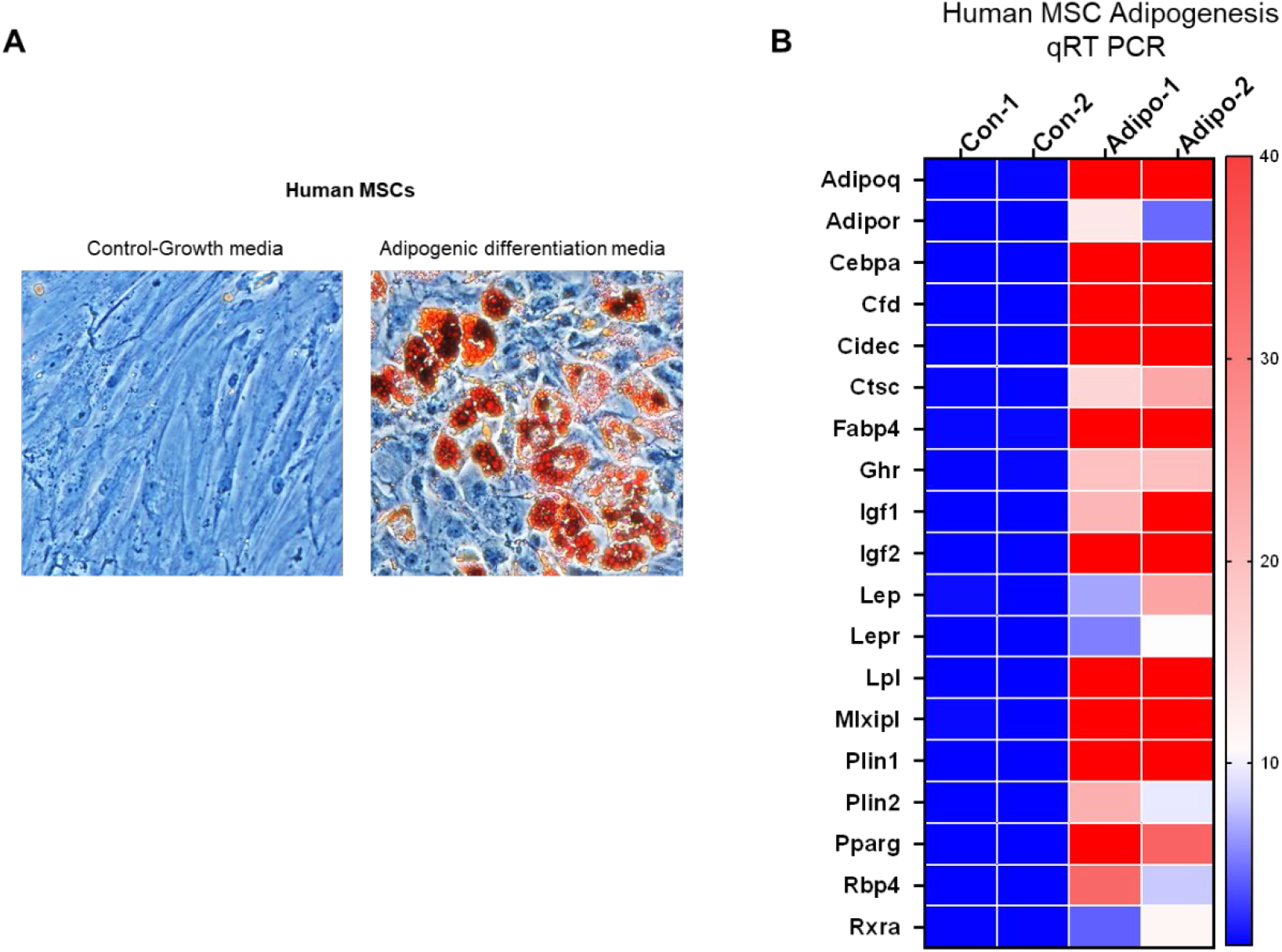
The genes which were linked by a STRING network, and selected for further validation *in vivo*, were validated by qRT-PCR in an adipocyte differentiation assay in human MSCs. mRNA from control and differentiated adipocytes were analyzed 2 weeks post-differentiation. The fold changes are shown as a heat map with fold changes above 40 represented as red.

**Supplementary Figure 2.**
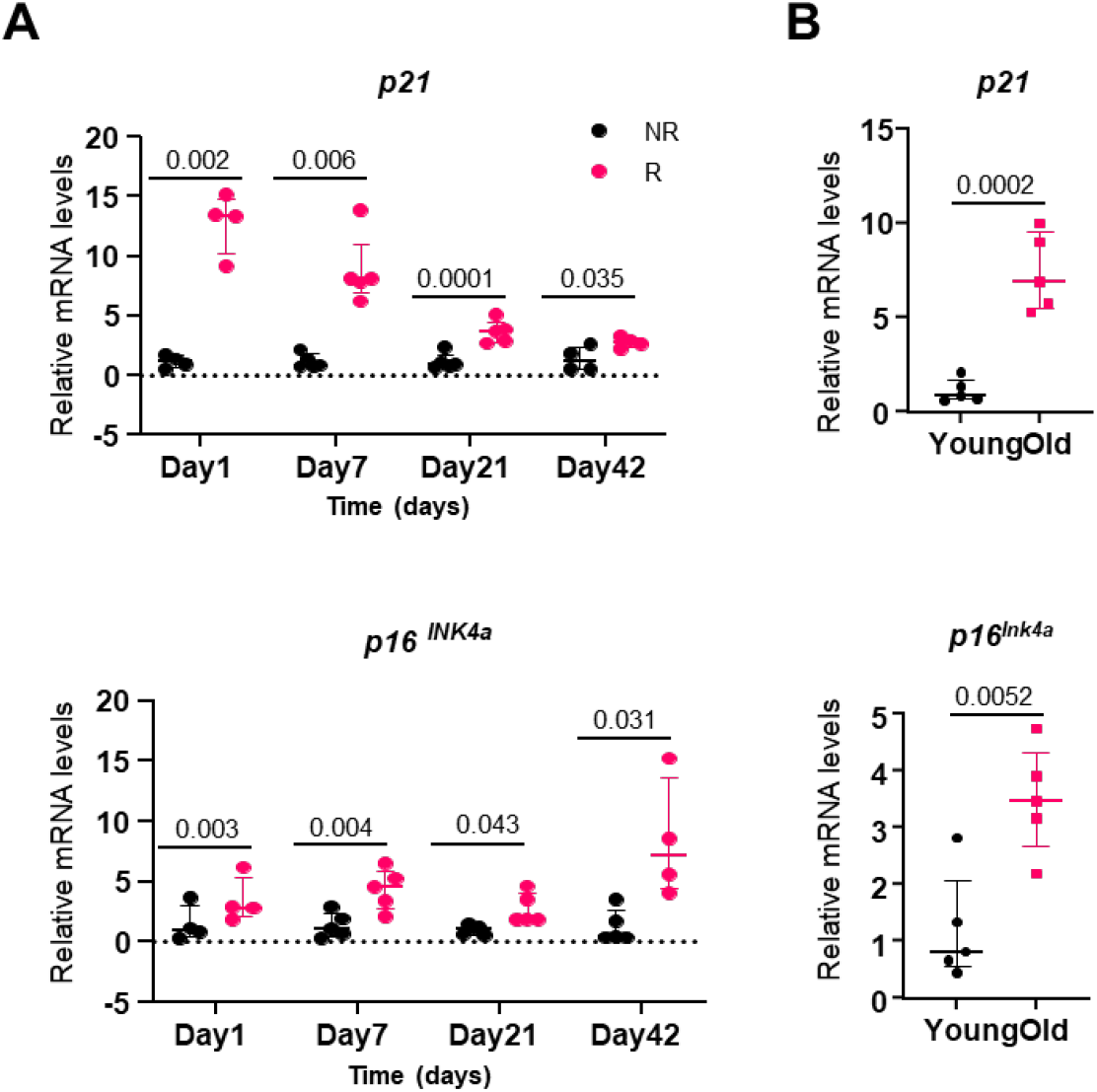
Comparative analysis of senescence markers in radiated and aged bones. Longitudinal assessments of senescence markers *Cdkn1a* (*p21*) and *Cdkn2a* (*p16*^*Ink4a*^) in R- and aged-bones. (A) The right femoral metaphyses of C57BL/6 mice were radiated (24Gy) in a 5 mm region, while the equivalent region of the left leg served as control. On days 1 (n=4),7 (n=5), 21(n=5) and 42 (n=4) R- and NR-femurs were harvested for mRNA isolation. Statistical analyses were done using GraphPad Prism and p-values were calculated using a two-tailed paired t-test. (B) Whole bones were collected from young (5m) (n=5) and old (24 m) (n=5) mice and mRNA were isolated. Results are expressed as medians with interquartile range. Statistical analyses were done using GraphPad Prism and p-values were calculated using a two-tailed unpaired t-test.

**Supplementary Table S1.**
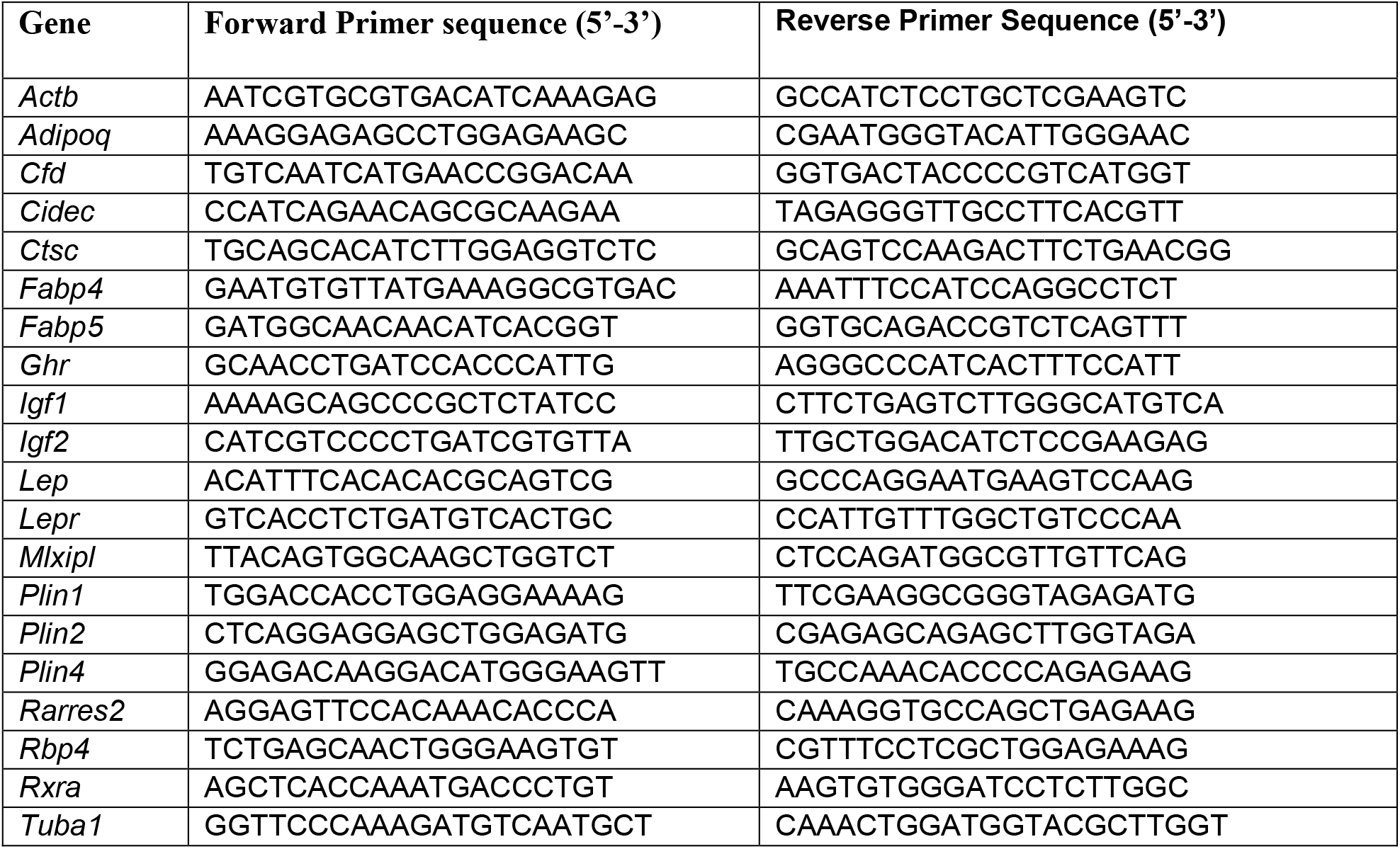
Mouse primer sequences

**Supplementary Table S2.**
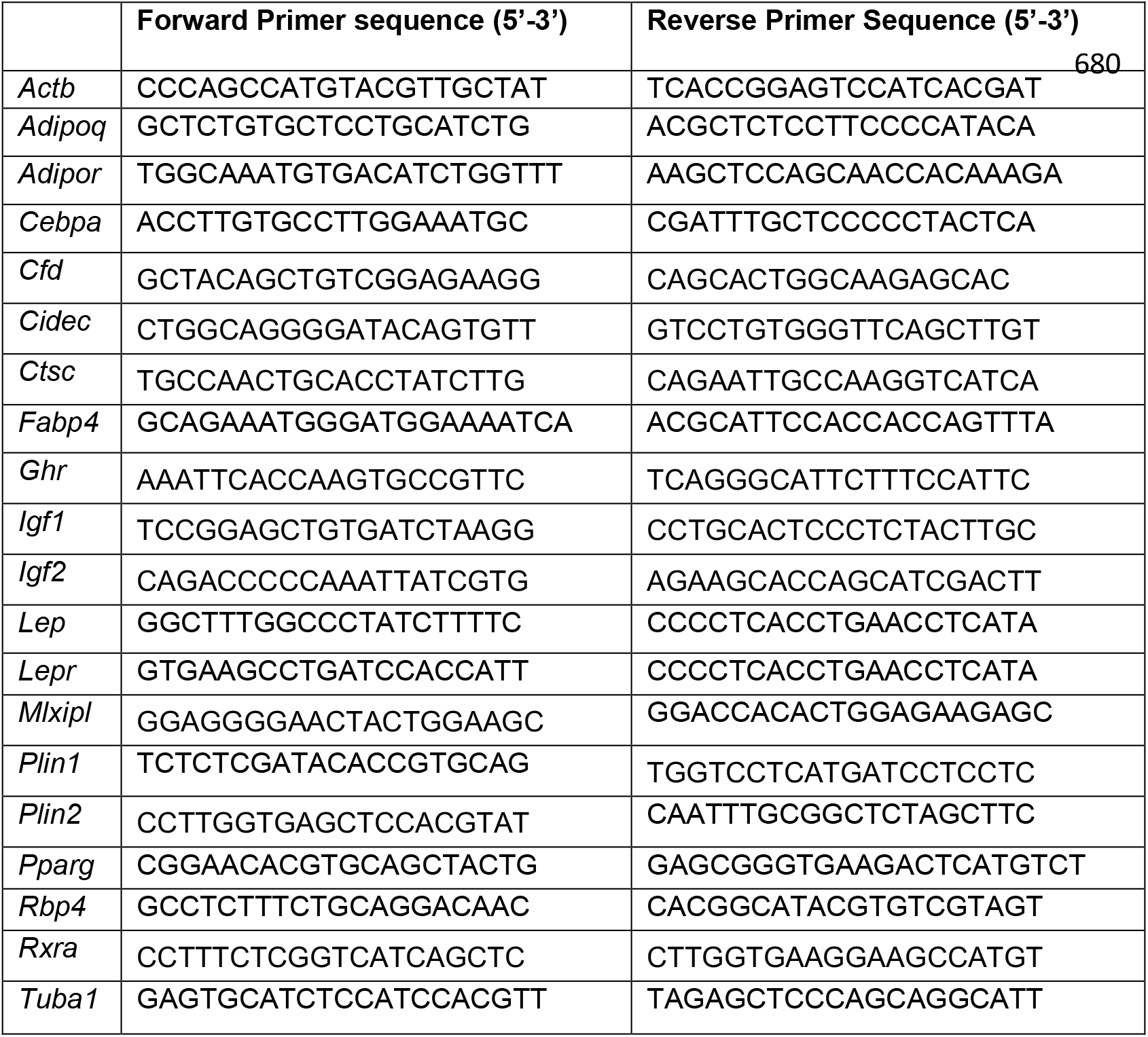
Human primer sequences

